# Cannflavin B ameliorates social and anxiety deficits and neuronal systems dysfunction in adolescent rats exposed to prenatal valproic acid

**DOI:** 10.1101/2025.09.14.676092

**Authors:** Olivia O. F. Williams, Madeleine Coppolino, Joshua D. Manduca, Taylor C. Demers, Paula T. Henry-Duru, Talen C. Mueller, Eric Soubeyrand, Colby J. Perrin, Tariq A. Akhtar, Melissa L. Perreault

**Author notes:** Corresponding Author: Olivia Williams, Department of Biomedical Sciences, 50 Stone Rd. E., University of Guelph, Guelph, Ontario, N1G 2W1, Canada.

## Abstract

There has been growing interest in natural products as potential therapeutics for the core and comorbid symptoms of autism spectrum disorders. Almost all the studies on autism have focused on the therapeutic benefits of cannabis and its associated cannabinoids. In this study the potential therapeutic efficacy of cannflavin B, a related, yet non-psychoactive component of the *Cannabis sativa* plant, was evaluated. Using prenatal valproic acid (VPA) exposure in rats, a model that has been widely used to study aspects of autism, we showed that cannflavin B was anxiolytic in the female VPA rats, and normalized sociality in VPA animals of both sexes. When neuronal oscillatory activity was examined, in female VPA rats cannflavin B normalized alterations in low frequency power within the cingulate cortex (Cg), and theta-gamma cross frequency coupling between the dorsal hippocampus (dHIP) and the prefrontal cortex (PFC). In male VPA animals, cannflavin B induced frequency-specific alterations in power within the PFC, Cg, and dHIP and ameliorated the VPA-induced suppression of oscillatory coherence between all three regions. In each brain region, cannflavin B also attenuated the sex-specific VPA-induced elevations in microglia. *In vitro*, cannflavin B normalized VPA-induced elevations in cortical and HIP neuronal activity and promoted more organized cortical firing. These findings demonstrate cannflavin B normalizes behavioural and neuronal systems function alterations induced by prenatal VPA in rats. The present study highlights the importance of alternative cannabis compounds in autism and other disorders.

## 1. Introduction

Autism spectrum disorders are a group of heterogeneous neurodevelopmental disorders that are diagnosed based on the expression of restricted and repetitive behaviours and difficulties socializing (American Psychiatric Association, 2013). Behavioural management or cognitive therapies are the predominant modes of treatment for autism, with pharmacological interventions primarily used to target the comorbidities, such as anxiety, aggression, and epilepsy (Matson and Goldin, 2013).

Cannabis and one of its major chemical constituents, cannabidiol (CBD), have been explored for their potential in treating the core symptoms of autism (da Silva et al., 2022; Riera et al., 2025). In autistic children, adolescents, or adults, whole cannabis extracts have been reported to improve sleep problems, attention deficit behaviours, repetitive behaviours, seizures, anxiety, as well as communication and personal interactions (Aran et al., 2019; David et al., 2024; Fleury-Teixeira et al., 2019; Montagner et al., 2023). Similarly, whole-plant cannabis extracts enriched with CBD improved communication and sociality scores in autistic children based on clinical and parental assessment (da Silva Junior et al., 2024; Hacohen et al., 2022). However, while findings are overall generally positive, there are also reports showing a lack of efficacy, worsening of symptoms, or the development of adverse effects (Aran et al., 2019; da Silva Junior et al., 2024; Hacohen et al., 2022; Montagner et al., 2023). Clinical studies examining the effects of CBD in autistic children also indicate improvements in some of their behavioural symptoms such as self-injurious behaviour, hyperactivity, sleep (Barchel et al., 2019), repetitive behaviour (Trauner et al., 2025), restlessness, aggression, anxiety, seizures, and agitation (Bar-Lev Schleider et al., 2019), as well as social impairments (Mazza et al., 2024). Similar to whole cannabis, some children experienced no improvements, whereas in others, symptoms worsened, or adverse side effects emerged (Barchel et al., 2019; Bar-Lev Schleider et al., 2019; Mazza et al., 2024; Trauner et al., 2025).

Although cannabis contains hundreds of chemical constituents, almost all the studies to mitigate ASD symptoms have focused on only two of the plant compounds, namely cannabidiol and tetrahydrocannabinol (THC). It has been suggested that several other compounds within the plant exhibit biological activity as well (Barrett et al., 1985; Werz et al., 2014). Flavonoids are a group of aromatic compounds that are found in numerous plant species and have been proposed as potential therapies in autism due to their antioxidant and anti-inflammatory properties (see review, Savino et al., 2023). In *Cannabis sativa*, the non-psychoactive flavonoids, cannflavin A and B, have exhibited cytoprotective and neuroprotective effects as well as antioxidant properties *in vitro* (Eggers et al., 2019; Li et al., 2023). Specifically, in PC12 cells, cannflavin A improved cell viability and protected against amyloid-β-induced neurotoxicity by reducing aggregate density and neurite loss (Eggers et al., 2019). In human keratinocyte cells, both cannflavin A or B were protective against erastin-induced cytotoxicity as a result of their antioxidant properties such as their ability to scavenge free radicals (Li et al., 2023). Cannflavin A and B have also demonstrated to have robust anti-inflammatory properties that are approximately 30 times more potent than aspirin by inhibiting microsomal prostaglandin E2 and leukotriene production (Barrett et al., 1985; Werz et al., 2014).

As neuroinflammation and oxidative stress have been strongly implicated in autism (Prata et al., 2019; Usui et al., 2023), in this study we therefore sought to identify whether cannflavin B could ameliorate the behavioural and neuronal systems function alterations inherent in rats prenatally exposed to valproic acid (VPA), a model widely used for the study of autism. Using *in vivo* and *in vitro* methodologies, the findings demonstrate that cannflavin B was highly efficacious in ameliorating the sex-specific behavioural and functional impacts of VPA.

## 2. Experimental Procedures

### 2.1 Microsomal production of cannflavin B

Cannflavin B was synthesized and purified using the methods of Holborn and colleagues (2023). The *Cannabis sativa* prenyltransferase CsPT3 recombinant protein was cloned and expressed in yeast (Holborn et al., 2023). Yeast cells expressing CsPT3 were re-suspended in 100 mM Tris-HCl, pH9.0, extracted with acid-washed glass beads (425 – 600 μM, Sigma-Aldrich) four times 30 second vortexed and kept on ice in between and centrifuged (1,500 × g, 20 minutes, 4 °C). The microsomes were purified from the supernatant by ultracentrifugation (160,000 × g, 90 minutes, 4 °C). The microsomes were re-suspended in 100 mM Tris-HCl, pH 9.0 and protein concentration determined by the Bradford method using a BSA standard curve. Enzymatic assays were performed for 4 hours at 37 °C with 1 mg/ml of microsomal protein, 200 μM of chrysoeriol and 400 μM of DMAPP in 100 mM Tris-HCl, pH 9.0 and 10 mM MgCl_2_. The reaction product was extracted by ethyl acetate phase partitioned twice and evaporated to dryness under N_2_ gas, and then re-suspended in methanol. The cannflavin B product was separated on a C18 reverse-phase column (Agilent Poroshell 120 EC-C18 – 150 mm × 4.6 mm, 2.7 μm) using a binary gradient with 45% methanol in water containing 0.1% formic acid (v/v); A) and 95% methanol in water containing 0.1% formic acid (v/v) (B), starting with 62% solvent B transitioning to 82% solvent B over the course of 20 minutes at a flow rate of 1 mL/min at 30 °C. Products were detected by absorption at 340 nm and quantified relative to authentic standards. Cannflavin B was collected off-line with a 6.2 minute retention time.

### 2.2 Animals

Pregnant Sprague-Dawley female rats (Charles River, QC) and their offspring were used. Rats were individually housed in large opaque rodent cages and kept on a 12-hour reverse light-dark cycle with food and water *ad libitum*. The valproic acid (VPA) model for the study of autism was employed for this work (Nicolini and Fahnestock, 2018). In humans, VPA exposure *in utero* often induces symptoms characteristic of autism (Bromley et al., 2013; Rasalam et al., 2005; Williams et al., 2001), and is therefore used to model learning, memory and sociality deficits that may mimic aspects of this group of disorders (Bertolino et al., 2017; de Mattos et al., 2020; Elesawy et al., 2022). Dams were administered a single intraperitoneal (i.p.) injection of VPA (500 mg/kg, concentration 250 mg/mL; Sigma-Aldrich Canada) or saline (SAL) (1 mL/kg), on gestational day 12.5 (Williams et al., 2025). For the *in vivo* studies, adolescent rat offspring were administered either 0.2 mg/kg cannflavin B (0.1 mg/mL dissolved in 5% DMSO in 0.9% saline) or equal volume vehicle (VEH). All protocols were in accordance with the guidelines set out by the Canadian Council on Animal Care Committee at the University of Guelph.

### 2.3 Behaviour

To evaluate anxiety, the elevated plus maze (EPM) was used. Animals were placed in the centre square of the maze, head facing an open arm, and allowed to explore for 5 minutes (Kraeuter et al., 2019). The total time spent in the open-arms, and number of entries into the open-arms were analyzed. Changes in social behaviour were studied using the three-chamber sociality test (Kim et al., 2014). The social apparatus consisted of three chambers, with the left and right chambers having a wire cage to temporarily house one animal for the duration of the experiment. Animals were habituated to the entire three-chamber social apparatus for 5 minutes immediately prior to testing. In the testing phase, the rat had access to the three chambers for 10 minutes, with the familiar and novel animals placed in either the left or right cage of the chamber. The familiar animal was defined as a cage-mate and the novel animal as an age- and sex-matched individual from a different litter. Familiar versus novel animal location was counterbalanced to eliminate location bias. Social index (SI) and social preference index (SPI) were measured as: SI = time spent socializing / time spent in empty chamber, and SPI = time spent interacting with the novel animal / time spent interacting with the familiar animal.

### 2.4 Surgeries

Electrode implantation surgeries were performed between postnatal day 36-38 as previously described (Albeely et al., 2024). Custom electrode microarrays were built using prefabricated Delrin templates and polyimide-insulated stainless-steel wires (A-M Systems: 791600, 0.008”). All arrays had an electrode impedance of less than 2MΩ. Bilateral stainless steel electrodes were implanted into the medial prefrontal cortex (PFC, AP +3.2, ML ±0.6, DV - 3.8), anterior cingulate cortex (Cg, AP +1.9, ML ±0.5, DV -2.8), and dorsal hippocampus (dHIP, AP -3.5, ML ±2.5, DV -2.5) (Paxinos and Watson, 2005). Animals were then single housed for recovery for a minimum of 7 days. Electrode placements were verified at the end of the study.

### 2.5 Local field potential recordings

Local field potential (LFP) recordings were taken from the PFC, the Cg, and dHIP (Albeely et al., 2024). Animals were habituated to the apparatus for two days for 5 minutes prior to recordings. All recordings (Wireless 2100-system, Multichannel Systems) were performed in awake and freely moving animals in clear plexiglass boxes (18″×18″×18″). Following baseline recordings animals were administered their allocated treatment with additional recordings taken immediately post-injection for 60 minutes at a sampling rate of 1000 samples/seconds. Spectral power and coherence between regions were analyzed using routines from the Chronux software package for MATLAB (MathWorks). Recordings were segmented, detrended, de-noised and low pass filtered to remove frequencies greater than 100 Hz. Continuous multitaper spectral power for the normalized data (to total spectral power) and coherence were evaluated in the following ranges for delta (1–4 Hz), theta (4–12 Hz), beta (12–32 Hz), and slow (32–60 Hz) and fast gamma (60 – 100 Hz). One minute time epochs were used for all spectral time courses and cross frequency coupling. Five minute epochs were taken at the end of the testing period for the coherence data.

### 2.6 Immunohistochemistry

Ninety minutes following the LFP recordings, animals were randomly selected from each group for the immunohistochemistry studies. Rats were perfused using 4% paraformaldehyde (PFA) and brain tissues were extracted, frozen, and stored at -80 °C. Fluorescence immunohistochemistry was performed on fixed free-floating brain sections (30 µm) (Albeely et al., 2022). Sections were first washed in 1X TBS (60.5 mM Tris, 87.6 mM NaCl pH 7.6), blocked for 2 hours (10% goat serum, 1% BSA, 0.2% Triton-X, 1X TBS), and incubated with mouse anti-Iba1 (1:200; Abcam, AB283319) primary antibody for 60 hours at 4 °C. Following incubation, sections were washed in 1X TBS, blocked (5% goat serum, 0.5% BSA, 0.01% Triton-X, 1X TBS), and incubated in an anti-mouse-Alexa 488 (1:200; Invitrogen) secondary antibody for 2 hours. Slices were then washed, and on the last wash 0.4 µg/mL of DAPI was added to stain nuclei (Cell Signaling, #4083). Brain slices were then mounted on slides using Prolong Gold (Thermo Fisher Scientific). Images were taken from the PFC, Cg, and dHIP at 20X magnification (Etaluma Lumascope).

### 2.7 Primary Neuronal Cell Cultures

Primary neuronal cortical and HIP cultures were generated from postnatal day 0-1 rat pups (Williams et al., 2025). The offspring were sexed and divided into four groups: female saline, male saline, female VPA, and male VPA. Frontal cortical and HIP tissue were dissected and placed in dissection media (water, 1X HBSS, 1% 1M HEPES, 1% penicillin/streptomycin) on ice. The tissue was then washed and dissociated in 0.5% Trypsin-EDTA at 37 °C for 20 minutes. Tissues were washed twice with dissection media, triturated, passed through a 100 μm pore cell strainer, and centrifuged at 200 x g for 5 minutes at room temperature. The supernatant was aspirated, and cells were resuspended in plating media (Neurobasal medium, 5% Fetal bovine serum, 2% B27 supplement, 1% 200 mM L-glutamine, and 1% penicillin/streptomycin). Cells were plated at a density of 3 x 10^5^ cells/well in 24-well CytoView multielectrode (MEA) plates (Axion Biosystems, Atlanta, GA, USA). Cell cultures underwent half-media changes with serum-free media (Neurobasal medium, 2% B27 supplement, 1% 200 mM L-glutamine, and 1% penicillin/streptomycin) two hours post-plating and maintained every 3-4 days for a total culturing period of 21 days.

### 2.8 Multielectrode array recordings

MEA recordings were performed on days *in vitro* (DIV) 21 using the Maestro Edge multi-well MEA recorder (Axion Biosystems, Atlanta, GA, USA) and Integrated Studio software (AxIS, Axion Biosystems, Atlanta, GA, USA) configured to neuronal-spontaneous activity (Williams et al., 2025). Cells were treated with 100 nM cannflavin B or DMSO, and recordings were taken 60 minutes post-treatment for 10 minutes at a sampling frequency of 12.5 kHz. The data were processed and analyzed using the NeuralMetric Tool software (Axion Biosystems). For the duration of the recording, plates were maintained at 37 °C and active electrodes were defined as >5 spikes per minute. The mean firing rate, number of bursts, inter-spike interval, and mean number of spikes/burst were extracted using data from all active electrodes within a treatment group. The synchrony index and coefficient of variation within a burst was calculated for each well with more than 5 active electrodes. Following spontaneous recording, extracellular waveform data were exported and analyzed with the Plexon Offline Sorter spike sorting tool (Offline Sorter v4.7.1, Plexon Inc., Dallas, TX, USA). Average peak, valley, and peak-valley distance were measured from each extracellular spike waveform that occurred over the 10 minute recording.

### 2.9 Statistical analyses

Prior to all statistical analyses, a Kolmogorov-Smirnov test (N ≥ 50) or Shapiro Wilk test (N ≤ 50) of normality was conducted to ensure data was equally distributed, and equality of variance assessed by Levene’s test. No data sets failed to meet normal distribution. A within sex univariate or repeated measures ANOVA was used. MEA data was collected from 16 electrodes/well for 3 wells from 3 biological replicates, for a total of N = 144 electrodes/group, electrodes were excluded if activity was < 5 spikes per minute (N = 98 – 144). Behavioural and *in vivo* electrophysiological data was collected from N = 10 – 12 rats/group. Immunofluorescence to detect Iba1 expression was performed N = 5 – 6 rats per group, with each hemisphere analyzed separately (N = 10 – 12 values/group). For quantification, the number of Iba1-positive cells were normalized to the total number of DAPI stained cells present in each slice, and each hemisphere measures were averaged from four slices. For all ANOVAs, group differences were assessed using Tukey or Games-Howell *post-hoc* tests as appropriate. All graphical data are presented as means ± SEM. Statistical significance is defined at *p* < 0.05.

## 3. Results

Cannflavin B was synthesized using microsomes expressing the *Cannabis sativa* prenyltransferase CsPT3 recombinant protein. The compound structure of cannflavin B is displayed in Figure 1A with the synthesized cannflavin B preparation, compared to an authentic standard, shown in Figure 1B. The experimental timeline is displayed in Figure 1C.

**Figure 1.**
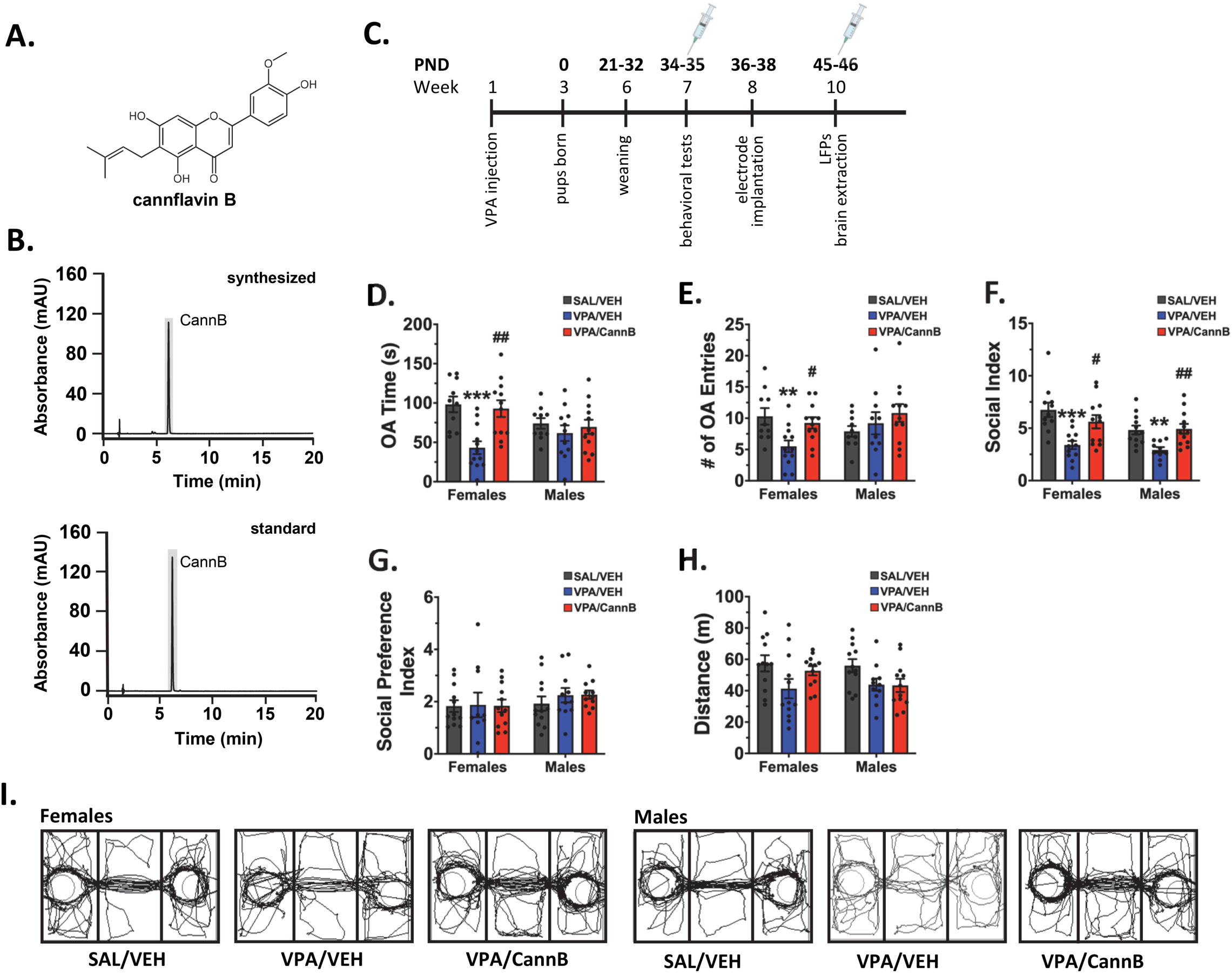
Cannflavin B ameliorated VPA-induced anxiety and social alterations. **A)** Molecular structure of cannflavin B, **B)** Chromatogram of isolated cannflavin B (CannB, upper panel) and the corresponding standard (lower panel), **C)** Experimental timeline, **D, E)** In the EPM, prenatal VPA exposure lowered the time spent in the open arms, and number of open arm entries for females only, effects normalized by cannflavin B (0.2 mg/kg), **F)** Prenatal VPA lowered the social index score in male and female rats, an effect lost with cannflavin B administration, **G, H)** No significant changes were observed in social preference index or total distance travelled, **I)** Representative tracings showing trajectories of movement in the three-chamber social apparatus. Data are expressed as means ± SEM. N = 10-12 animals/group. ** *p* < 0.001, *** *p* < 0.001 compared to SAL/VEH controls, ^#^ *p* < 0.05, ^##^ *p* < 0.01 compared to VPA/VEH group, Tukey or Games-Howell post-hoc test.

### 3.1 Cannflavin B is anxiolytic and increases sociability

The effect of cannflavin B on VPA-induced anxiety was first evaluated in adolescent rats. In the EPM, only the female VPA rats exhibited elevated anxiety-like symptoms as indicated by the reduced time spent in the open arm (*p* < 0.001) and reduced number of open arm entries (*p* = 0.009, Fig. 1D, E), and these measures were normalized following cannflavin B administration. [open arm time: F(2,31) = 10.1, *p* < 0.001; # of open arm entries: F(2,31) = 5.8, *p* = 0.007]. No group differences in anxiety were observed in the male animals.

The social index (SI) is a measure of the total time the rodents spend interacting with both the novel and familiar animals compared to the empty chamber, with a higher SI being an indicator of greater socialization. Both female and male VPA animals were less social compared to their sex-matched controls (*p* < 0.001, *p* = 0.009 respectively), and these effects were normalized by cannflavin B (Fig. 1F). There were no significant group differences observed in the preference between a novel and a familiar rat (Fig. 1G). Evaluating overall locomotor activity in the social apparatus showed no effects of Cannflavin B (Fig. 1H). Representative tracings of animals in the three-chamber social apparatus are identified in Figure 1I. [SI: Female, F(2,32) = 8.7, *p* < 0.001; Male, F(2,31) = 6.9, *p* = 0.003].

### 3.2 Cannflavin B ameliorates neuronal oscillatory activity alterations

Neuronal oscillations are highly coupled to behavioural and disordered states (Uhlhaas and Singer, 2010; Zikopoulos and Barbas, 2013) and have been suggested as potential biomarkers of therapeutic efficacy (Arutiunian et al., 2024; Gandal et al., 2010). We therefore evaluated the effects of cannflavin B treatment on neuronal oscillatory activity in, and between, the PFC, Cg, and dHIP (Figs. 2 and 3). Regions of electrode placements are shown in Figure 2A. For the female rats, power spectra for each region are depicted in Figure 2B-D, with the time courses across the 60 minute recording period shown in Figure 2E-G. VPA exposed female rats exhibited frequency-specific alterations in spectral power selectively in the Cg, with repeated measures analysis showing an overall lower delta (*p* = 0.034), and higher theta (*p* = 0.035) and beta (*p* = 0.030) power across the entire recording (Fig. 2F). These alterations were ameliorated by cannflavin B administration. There were no group differences in inter-regional coherence between any of the brain regions for the female animals (Fig. 2H-J). Cross-correlation analysis revealed a reduction in the strength of phase coupling between dHIP theta and low gamma in the PFC, but not the Cg, of VPA-exposed female rats, which were sensitive to the effects of cannflavin B (Fig. 2K and Fig S1A). Female VPA rats also exhibited a 30 ms delay in dHIP-PFC theta-low gamma coupling that was normalized following administration of cannflavin B (Fig. 2K).

**Figure 2.**
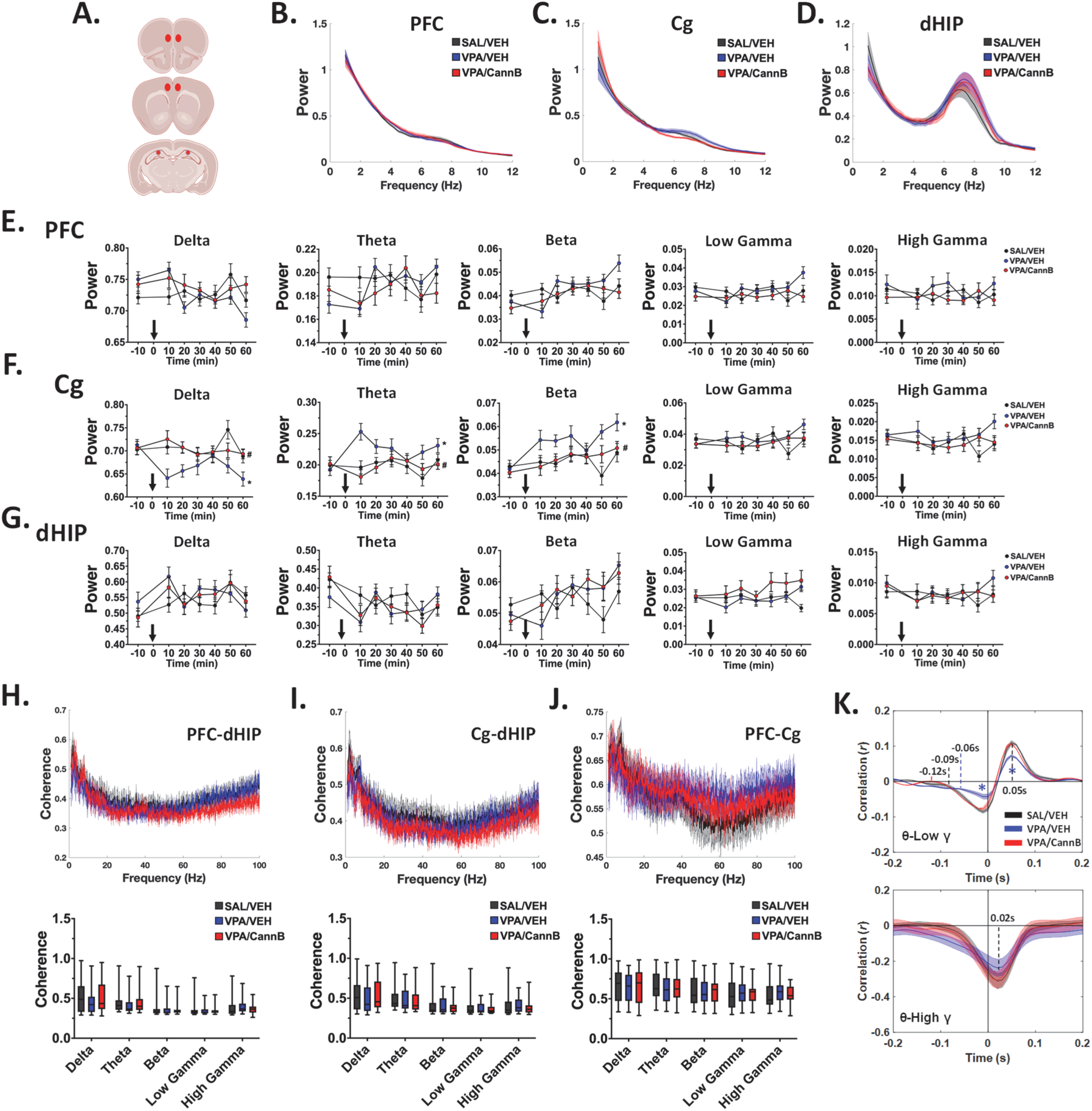
Effects of cannflavin B on prenatal VPA-induced alterations in neuronal oscillatory activity in female rats. **A)** Areas of electrode placement, **B-D)** Representative power spectra for the PFC, Cg, and dHIP, **E-G)** Time courses showing effects of VPA and cannflavin B (CannB) treatment (0.2 mg/kg) on spectral power for each frequency across the 60 min recording period. Data shown at each time point are derived from 1 min epochs, **H-J)** No effect of VPA alone or following cannflavin B on inter-regional coherence, **K)** Alterations in dHIP-PFC theta-gamma coupling in female rats exposed to prenatal VPA alone and with cannflavin B administration. Data are expressed as means ± SEM. N=10-12 animals/group. Time courses were analyzed by repeated measures ANOVA. * *p* < 0.05, compared to SAL/VEH controls, ^#^ *p* < 0.05, compared to VPA/VEH group, Tukey or Games-Howell post hoc.

**Figure 3.**
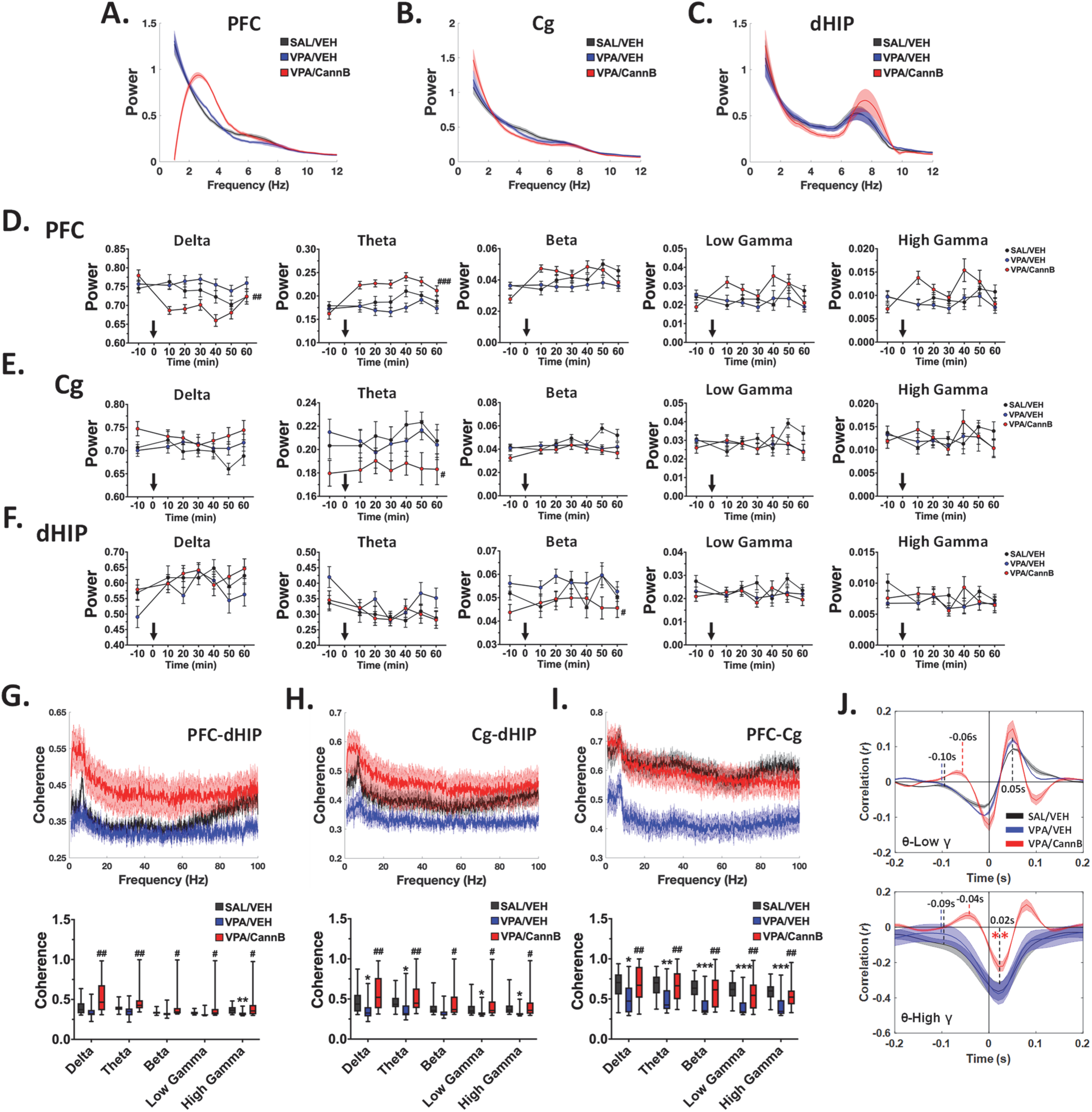
Effects of cannflavin B on prenatal VPA-induced alterations in neuronal oscillatory activity in male rats. **A-C)** Representative power spectra for the PFC, Cg, and dHIP, **D-F)** Time courses showing effects of VPA and cannflavin B (CannB) treatment (0.2 mg/kg) on spectral power for each frequency across the 60 min recording period. Data shown at each time point are derived from 1 min epochs, **G-I)** Prenatal VPA suppressed inter-regional coherence across frequencies and this was normalized by cannflavin B, **J)** Cannflavin B induced timing alterations in dHIP-PFC theta-low gamma coupling and reduced the strength of dHIP-PFC theta-high gamma coupling in male rats exposed to prenatal VPA. Data are expressed as means ± SEM. N=10-12 animals/group. Time courses were analyzed by repeated measures ANOVA. ^#^ *p* < 0.05, ^##^ *p* < 0.01, ^###^ *p* < 0.001 compared to VPA/VEH group, Tukey or Games-Howell post hoc.

Representative power spectra for each region in male rats are depicted in Figure 3A-C, with the brain region-specific time courses shown in Figure 3D-F. There were no VPA-induced alterations in spectral power in any of the regions evaluated, though some effects were observed in VPA rats treated with cannflavin B. Specifically, in the PFC, cannflavin B suppressed delta (*p* = 0.001) and elevated theta (*p* < 0.001) power in the VPA animals (Fig. 3D). In the Cg and dHIP cannflavin B induced a significant, though modest, suppression of theta (*p* = 0.042) and beta (*p* = 0.017) power respectively (Fig. 3E, F). Prominent reductions in inter-regional coherence were observed in the VPA-exposed male animals (Figs. 3G-I). Lower dHIP-PFC gamma coherence (*p* = 0.001) was evident, as well as lower dHIP-Cg delta (*p* = 0.043), theta *(p* = 0.033), and low and high gamma (*p* = 0.037 and *p* = 0.018 respectively) (Fig. 3G, H). A frequency-wide suppression of PFC-Cg coherence was also apparent (delta: *p* = 0.028; theta: *p* = 0.002; beta: *p* < 0.001; low gamma: *p* < 0.001; high gamma: *p* < 0.001; Fig. 3I). All the suppressive effects on coherence in the VPA animals were ameliorated by cannflavin B administration.

Analysis of cross-frequency coupling in the male rats showed significant treatment effects on theta-gamma coupling between the dHIP and PFC (Fig. 3J). There were no differences in theta-low gamma coupling strength with VPA exposure alone or in combination with cannflavin B. However, cannflavin B altered the timing of dHIP theta coupling to both the low and high gamma frequencies in the PFC by 40-50 milliseconds and significantly altered the time-dependent theta-high gamma coupling strength (Fig. 3J). There was a minor 10 millisecond shift in the timing of dHIP-Cg theta-low gamma coupling with no group differences in dHIP-Cg theta-high gamma coupling (Fig. S1B).

### 3.3 Attenuation of VPA-induced microglial activity by cannflavin B

The impact of cannflavin B on VPA-induced alterations in microglial activation was next evaluated. Representative images of Iba1-positive microglia for each group in each of the brain regions are shown (Fig. 4A). In the PFC, female VPA animals had elevated Iba1-positive cells compared to controls (*p* = 0.035, Fig. 4B) that was attenuated by cannflavin B [PFC: F(2, 33) = 9.9, *p* < 0.001]. Though VPA exposure did not alter basal microglial activation in the PFC of male animals, a significant reduction in microglial activity by cannflavin B in male VPA rats was evident (*p* < 0.001, Fig. 4B). In the Cg and dHIP, only the male VPA animals showed elevated levels of Iba1-positive cells (*p* = 0.002 and *p* = 0.004 respectively, Fig. 4C, D), effects that were diminished by cannflavin B. [PFC: F(2, 31) = 9.5, *p* < 0.001; Cg: F(2, 31) = 9.7, *p* < 0.001; dHIP: F(2, 31) = 9.5, *p* < 0.001].

**Figure 4.**
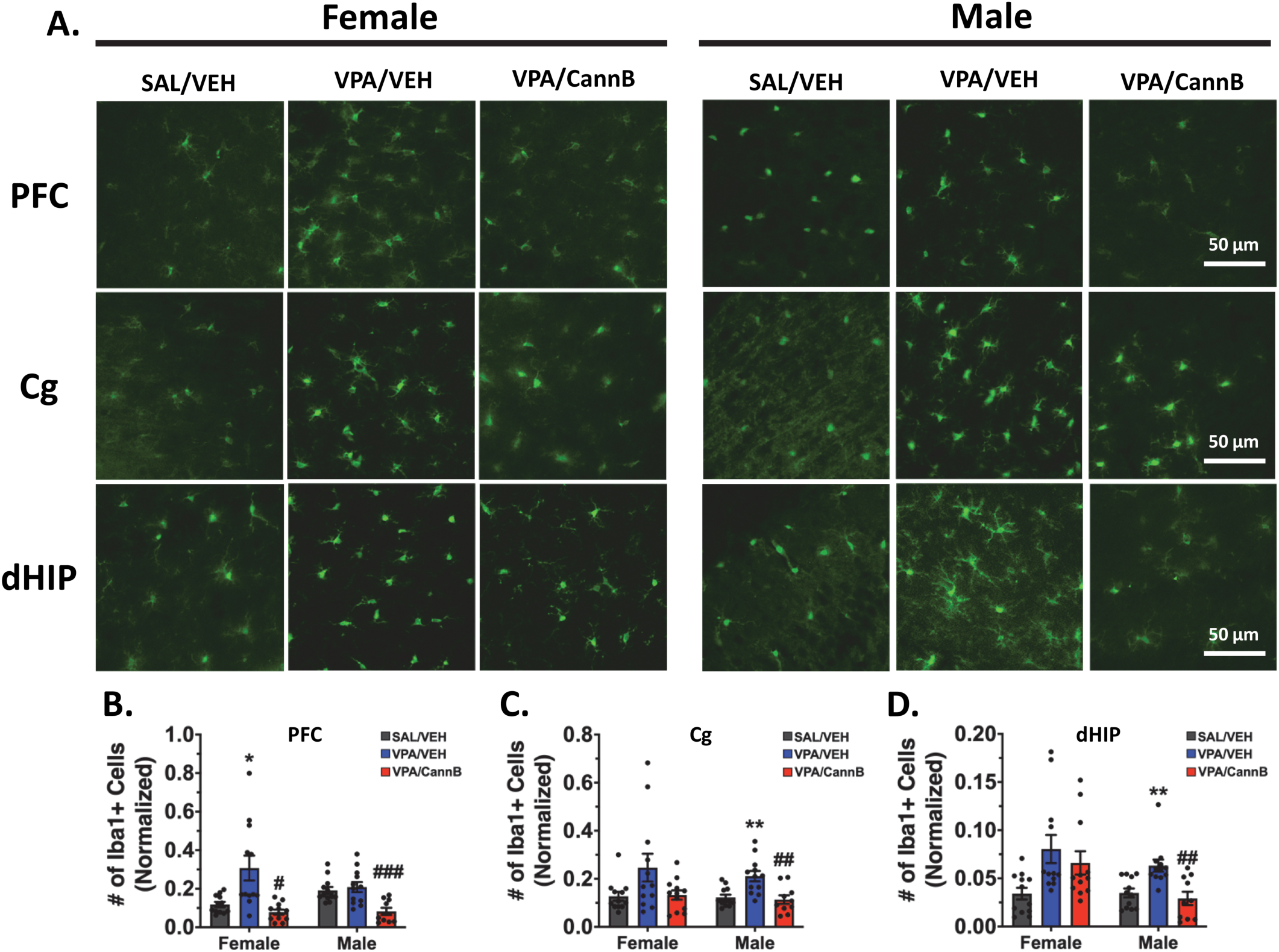
Attenuation of prenatal VPA-induced elevations in microglial activation in PFC, Cg, and dHIP by cannflavin B. **A)** Representative images of Iba1+ cells in male and female rats exposed to prenatal VPA and cannflavin B (CannB, 0.2 mg/kg) effects on expression in the PFC, Cg, and dHIP, **B-D)** Quantification of Iba1-expressing cells in each of the three brain regions. VPA-induced elevations in microglial activity were suppressed by cannflavin B. Data are expressed as means ± SEM. N = 5-6 animals/group, 2 slices/rat with hemispheres averaged. Iba1+ cells were normalized to the number of cells present in each image (for clarity, DAPI not shown). \**p* < 0.05, and \*\**p* < 0.01 compared to VPA/VEH group, ^##^ *p* < 0.01, ^###^ *p* < 0.001 compared to VPA/VEH group, Tukey or Games-Howell post hoc.

### 3.4 Cannflavin B normalizes aberrant neuronal firing *in vitro*

We have previously shown sex-specific alterations in neuronal activity in cortical and HIP primary neurons derived from rodents prenatally exposed to VPA (Williams et al., 2025). We therefore sought to determine whether cannflavin B could ameliorate these effects. Raster plots for each sex showing group differences in neuronal activity are shown (Fig. 5A). Both female and male VPA cortical neurons had elevated activity compared to sex matched controls as observed by the mean firing rate (female, *p* < 0.001; male, *p* < 0.001), and number of spikes per burst (female, *p* = 0.011; male, *p* < 0.001) (Fig. 5B, D). Female cortical VPA neurons additionally displayed elevated bursting activity (*p* < 0.001) and a higher firing synchrony (*p* < 0.001) (Fig. 5C, E), as well as a lower inter-spike interval (*p* < 0.001) (Fig. 5F). Cannflavin B attenuated VPA-induced elevations in mean firing rate for both sexes, and number of bursts for female VPA neurons only (Fig. 5B, C). Despite no previous VPA-induced change, cannflavin B significantly reduced the number of bursts in male VPA neurons (*p* < 0.001) (Fig. 5C). Both female and male cortical VPA neurons also had more disorganized firing (CV: female, *p* < 0.001; male, *p* < 0.001) which was normalized by cannflavin B selectively in the female VPA neurons (Fig. 5G, H). The proportion of extracellular waveform types (regular, fast, positive, compound, triphasic) did not differ between groups for either sex. However, cortical VPA neurons from each sex had overall lower amplitudes (peak: female, *p* = 0.023; male, *p* = 0.046; valley: female, *p* = 0.001; male, *p* = 0.038) (Fig. 5I-K), as well as a shorter peak to valley distance (female, *p* = 0.007; male, *p* = 0.035) (Fig. 5I, L), indicating that the neurons had a lower depolarization threshold and recovered more rapidly. These alterations were not affected by cannflavin B treatment.

**Figure 5.**
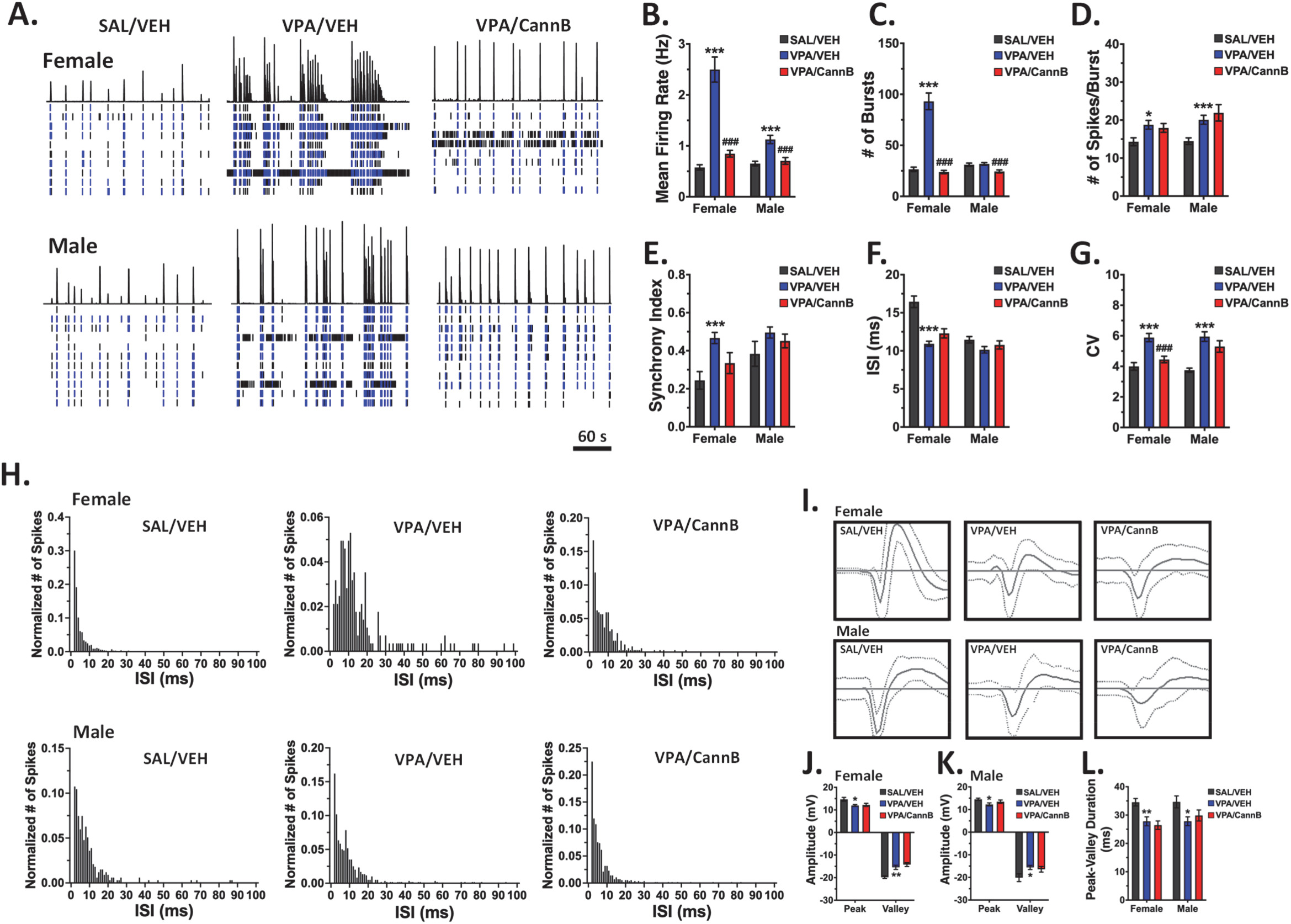
Effects of cannflavin B on neuronal activity of cortical neurons *in vitro* derived from rats prenatally exposed to VPA. **A)** Representative raster plots from male and female derived cortical neurons showing group differences in neuronal spiking, **B, C)** Elevations in the mean firing rate in VPA neurons of both sexes, and bursting in the female VPA neurons, was normalized by cannflavin B (100 nM) treatment, **D)** VPA elevated the number of spikes/burst with no cannflavin B effects, **E-F)** Cannflavin B reduced VPA-induced elevations in synchronous activity in female-derived neurons, with no effect on interspike interval (ISI), **G)** Female and male VPA neurons had more disorganized firing, indicated by a higher coefficient of variation (CV), which was normalized by cannflavin B in the females only, **H)** Representative ISI histograms for each experimental condition, **I-K)** Representative extracellular waveforms and quantification showing a smaller peak and valley amplitude in female and male VPA neurons with no effects of cannflavin B, **L)** Peak-valley duration was suppressed in the VPA neurons with no effects of cannflavin B. Data are expressed as means ± SEM. N=3 wells/treatment from 3 biological replicates. * *p* < 0.05, ** *p* < 0.01, *** *p* < 0.001 compared to VPA/VEH group, ^###^ *p* < 0.001 compared to VPA/VEH group, Tukey or Games-Howell post hoc.

In the HIP cultures, female but not male VPA neurons had a higher mean firing rate (*p* < 0.001) and this was normalized by cannflavin B (Fig. 6A, B). Conversely, male but not female HIP VPA neurons exhibited lower bursting activity compared to sex-matched control neurons (*p* < 0.001), with no improvement by cannflavin B (Fig. 6A, C). The number of spikes per burst in the VPA neurons were also different between the sexes, with a lower number of spikes per burst in the female VPA neurons (*p* = 0.014) and a higher number for the male VPA neurons (*p* = 0.006) (Fig. 6D). However, cannflavin B ameliorated the effects of VPA selectively in the male HIP neurons (Fig. 6D). Only the male VPA neurons had a lower synchrony index (*p* = 0.002) and a higher inter-spike interval (*p* < 0.001), neither of which were improved with cannflavin B treatment (Fig. 6E, F). Cannflavin B promoted more organized firing in the female VPA neurons (*p* < 0.001) (Fig. 6G, H). Similar to that observed with the cortical neurons, there were no group differences in the proportion of waveform types observed. There were no sex or group differences in waveform amplitude or peak-valley duration (Fig. 6I-L).

**Figure 6.**
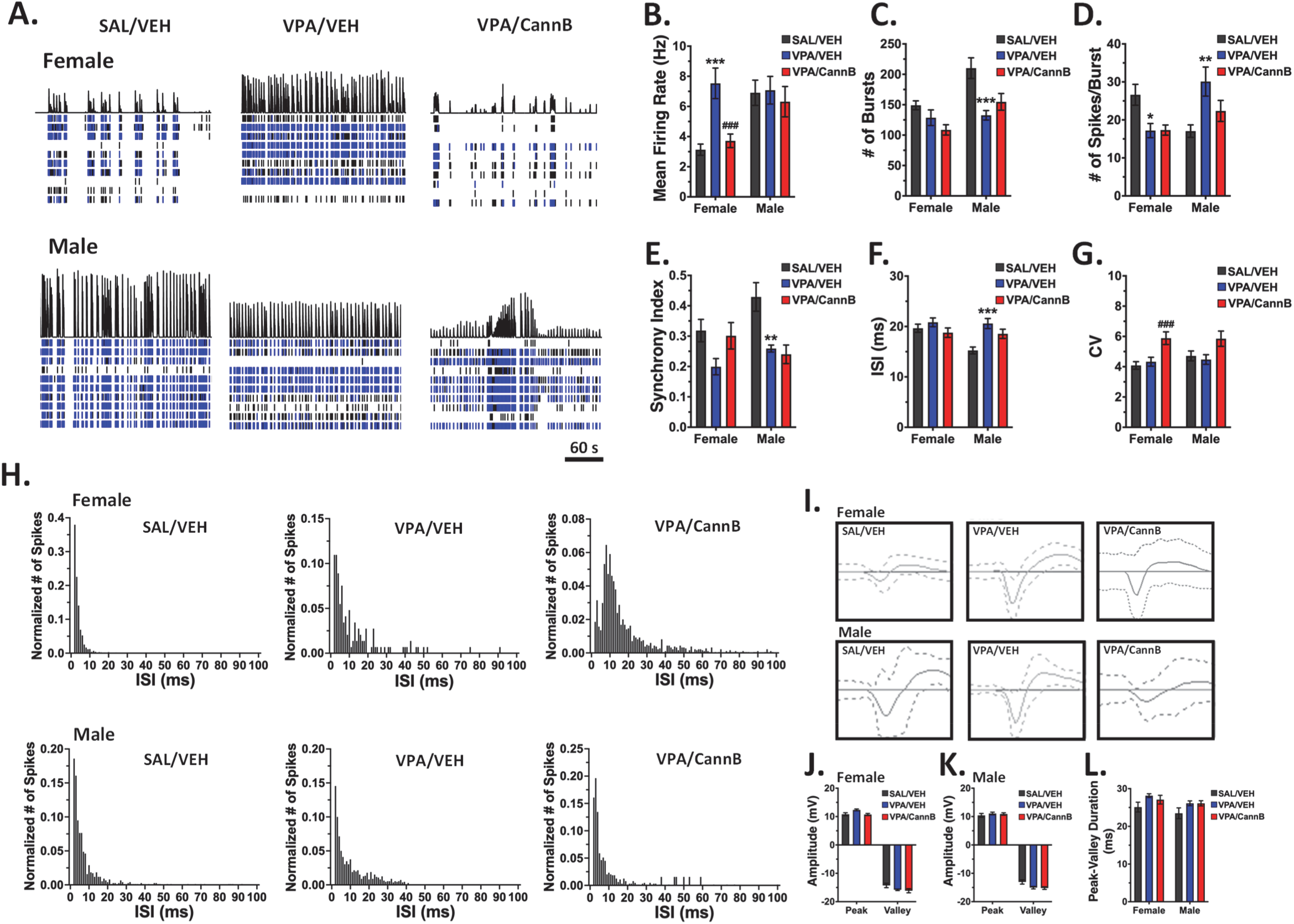
Effects of cannflavin B on neuronal activity of HIP neurons *in vitro* derived from rats prenatally exposed to VPA. **A)** Representative raster plots from male and female derived HIP neurons showing group differences in neuronal spiking, **B)** VPA elevated the mean firing rate in female-derived neurons that was attenuated by cannflavin B (100 nM), **C-F)** Sex-specific alterations in bursting activity, the number of spikes/burst, synchrony, and the interspike interval (ISI) induced by VPA were not altered by cannflavin B, **G)** No innate effect of VPA on the coefficient of variation (CV) in HIP neurons, though cannflavin B elevated the CV in female VPA neurons, **H)** Representative ISI histograms for each experimental condition, **I-L)** No effect of VPA or cannflavin B on HIP neuron extracellular waveform dynamics. Data are expressed as means ± SEM. N=3 wells/treatment from 3 biological replicates. * *p* < 0.05, ** *p* < 0.01, *** *p* < 0.001 compared to VPA/VEH group, ^###^ *p* < 0.001 compared to VPA/VEH group, Tukey or Games-Howell post hoc.

## 4. Discussion

In the present study, we provide the first *in vivo* neurobiological characterization of cannflavin B, a non-psychotropic flavonoid found in the *Cannabis sativa* plant. Rats prenatally exposed to VPA manifest sex-specific behavioural and neurobiological changes, and we showed that many of these alterations were ameliorated by cannflavin B. Acute cannflavin administration to VPA-exposed adolescent rats improved socialization in both sexes and was anxiolytic in the female animals. Cannflavin B also normalized VPA-induced alterations in neuronal oscillatory activity and inflammatory responses. These findings were supported by the *in vitro* studies in cortical and HIP neurons that showed cannflavin B could ameliorate most of the neuronal activity changes induced by VPA.

There are few studies to date that have examined the neurobiological effects of cannflavin B, though some evidence indicates the compound has potent anti-inflammatory activity (Barrett et al., 1985; Werz et al., 2014). These anti-inflammatory properties were shown to be mediated through its inhibitory effects on prostaglandin E2 (PGE2) signaling (Barrett et al., 1985; Werz et al., 2014) via suppression of microsomal prostaglandin E synthase-1 (mPGES-1) (Abdel-Kader et al., 2023; Werz et al., 2014). PGE2 is primarily produced through the conversion of arachidonic acid by cyclooxygenases (COX) enzymes, COX-1 and COX-2, through a multi-step process involving mPGES-1 (Chen et al., 2022; Jiang et al., 2017). COX-1 is constitutively expressed and is involved in maintaining homeostatic functions, whereas COX-2 is inducible in response to a variety of pro-inflammatory stimuli (Faki and Er, 2021). Both enzymes are targets of non-steroidal anti-inflammatory drugs, though some drugs such as aspirin show greater affinity for COX-1. Further, mPGES-1 functions downstream of the COX enzymes to convert prostaglandin H2 to PGE2, which can subsequently bind to its various receptors (EP1, EP2, EP3, and EP4). The prostaglandin receptors have been demonstrated to be involved in both neurotransmitter release and synaptic plasticity (Akaneya, 2008; Li et al., 2018), as well as in inflammation (Jiang and Dingledine, 2013). Alterations in levels of prostaglandin and COX-2 have previously been reported in autism (El-Ansary et al., 2016; El-Ansary and Al-Ayadhi, 2012; Qasem et al., 2018). Reports indicate that individuals diagnosed with autism have elevated blood PGE2, mPGES-1, and COX2 levels (El-Ansary and Al-Ayadhi, 2012; Qasem et al., 2018), with changes in mPGES-1 as one of the best predictors of the degree of sensory profile impairment (El-Ansary et al., 2016; Qasem et al., 2018). It has been suggested that alterations in COX-2 activity and prostaglandin signaling are the core mediators of enhanced neuroinflammation in autism, and may also be involved in regulating disruptions in synaptic plasticity (Kamal et al., 2025). Inhibition of mPGES-1 by cannflavin B could therefore have therapeutic properties in this group of disorders by suppressing the heightened PGE2 levels that are driven by elevated activity of the COX or mPGES-1 enzymes.

Cannflavin B also inhibits 5-lipoxygenase (Werz et al., 2014), an enzyme involved in the conversion of arachidonic acid into leukotrienes. There is less know about the role of leukotrienes in autism, though some evidence of increased leukotriene levels have been reported (El-Ansary and Al-Ayadhi, 2012). Furthermore, in a study utilizing the Pilo mouse model of epilepsy, which displays autism-like features, gene deletion of 5-lipoxygenase improved learning and memory, and socialization (Guan et al., 2024).

Flavonoids in general have been recognized for their antioxidant properties (Bjørklund et al., 2020; Liu et al., 2022; Savino et al., 2023). For example, flavonoids have been reported to inhibit inducible nitric oxide synthase levels, scavenge reactive oxygen species, as well as increase glutathione to prevent oxidative stress (Bertolino et al., 2017; de Mattos et al., 2020; Elesawy et al., 2022; Li et al., 2023). Indeed, both cannflavin A or B were similarly shown to scavenge free radicals in human keratinocytes (Li et al., 2023). Some studies have demonstrated beneficial effects of flavonoid treatment in regulating behaviour in the VPA model (Bertolino et al., 2017; de Mattos et al., 2020; Elesawy et al., 2022) as well as in humans diagnosed with autism (Bertolino et al., 2017), effects that were associated to the flavonoids’ antioxidant properties. Other translational work also implicates flavonoid-induced alterations in neurotransmitter function as being potentially beneficial in autism (Serra et al., 2022).

*In vitro* findings in primary cortical neurons showed that cannflavin B has antagonist properties at the tropomyosin receptor kinase B (TrkB), attenuating brain-derived neurotrophic factor (BDNF)-induced accumulation of activity-regulated cytoskeleton-associated (Arc) protein, and Akt and mTOR phosphorylation (Holborn et al., 2023). In autism, BDNF expression can be upregulated (Almeida et al., 2014; Barbosa et al., 2020; Connolly et al., 2006; Correia et al., 2010; Nishimura et al., 2007). For example, elevated BDNF concentrations in serum (Barbosa et al., 2020; Connolly et al., 2006) and platelet rich plasma (Correia et al., 2010) from autistic children were observed. Elevated BDNF mRNA was also reported in lymphocytes from blood samples of drug-naïve autistic individuals (Nishimura et al., 2007), an observation also found in fetal brain tissue post VPA administration *in utero* (Almeida et al., 2014).

A main protein regulated by TrkB activity is glycogen synthase kinase-3 (GSK-3). Phosphorylated by the downstream effector of TrkB, Akt, to suppress its activity, GSK-3 mediates an array of physiological functions including mediating intracellular signaling, as well as regulating neuronal plasticity, gene expression, and cell survival (Grimes and Jope, 2001). GSK-3 is also a key downstream effector of Wnt signaling, a pathway that is integral during the developmental period and implicated in various neurodevelopmental disorders including autism (Mulligan and Cheyette, 2017). Although GSK-3 has been well-studied for its role in cognition and its involvement as a critical mediator of cognitive decline across numerous disease states (Ances et al., 2008; King et al., 2013; Takashima, 2006), a role for the kinase in autism is emerging. In children with autism, reduced activity of GSK-3 in isolated T cells has been reported (Onore et al., 2017). Consistent with this finding, in the VPA model a similar decrease in GSK-3 activity, with an associated increase in the activity of its effector β-catenin, was shown in various brain regions including PFC, HIP, and cerebellum (Qin et al., 2016), as well as amygdala (Wu et al., 2017). Activation of the GSK-3/β-catenin pathway was also shown to underlie macrocephaly in VPA animals (Go et al., 2012), a phenomenon also observed in autistic individuals (Landa and Garrett-Mayer, 2006; Wetherby et al., 2004).

Neuroimaging studies to date have defined a putative brain network mediating symptoms in autism (Ecker et al., 2017) with the most replicable findings involving the cortical and limbic regions. In line with these studies, disturbances in neural oscillatory activity and functional connectivity in the network have also been demonstrated (Kessler et al., 2016). For example, it has been suggested that a cortical inhibitory/excitatory imbalance, induced through dysfunction of local inhibitory neurons, increases local synaptic connectivity and manifests as increased gamma frequency or “noisy” activity in local circuits (Berg and Plioplys, 2012; Spence and Schneider, 2009). These alterations would also have impacts on long-distance communication resulting in aberrant connectivity to other cortical regions (Casanova et al., 2002; Courchesne and Pierce, 2005; Lewis and Elman, 2008). There is also accumulating evidence of low frequency deficits in delta/theta long-range coupling (Kessler et al., 2016) that could potentially play a role in the known dysregulation in connectivity within networks involving the thalamus, striatum, and amygdala (Cerliani et al., 2015; Glerean et al., 2016; Kleinhans et al., 2016). Here we report significant sex-specific alterations in oscillatory activity in VPA rats, including region-specific alterations in low frequency oscillatory power, a dysregulation of HIP-PFC theta-gamma coupling, and suppressed oscillatory coherence in male VPA rats, with most of these alterations normalized or diminished following cannflavin B administration. Of relevance to the present work, our past findings in male rats have linked changes in PFC or HIP GSK-3 activity directly to alterations in oscillatory activity and learning and memory dysfunction, and it was proposed that GSK-3 may play a homeostatic role in brain systems function with aberrant up or downregulation in activity having negative effects on cognitive performance (Albeely et al., 2022). As evidence suggests GSK-3 activity is downregulated following prenatal VPA exposure (Williams et al., 2025), it is therefore possible that the normalization of GSK-3 activity by cannflavin B was involved in the observed improvements in brain systems function reported herein.

In conclusion, this was the first study to examine the *in vivo* effects of cannflavin B in a model widely used to study aspects of autism. It was demonstrated that cannflavin B, a non-psychoactive component of the *Cannabis sativa* plant, could normalize many of the sex dependent and independent alterations in behaviour and neuronal activity, as well as microglial activation, in adolescent rats following prenatal exposure to VPA. Though still at the early stages of investigation, the present findings provide evidence of significant potential for cannflavin B to ameliorate the social impairments and anxiety symptoms of autism. In depth learning and memory studies should be performed, as well as research delineating the specific mechanisms that underlie these effects, potentially focusing on the involvement of PGE2 and/or TrkB signaling and downstream mediators.

