## Supplemental Figure 1 for "Cannflavin B ameliorates social and anxiety deficits and neuronal systems dysfunction in adolescent rats exposed to prenatal valproic acid"

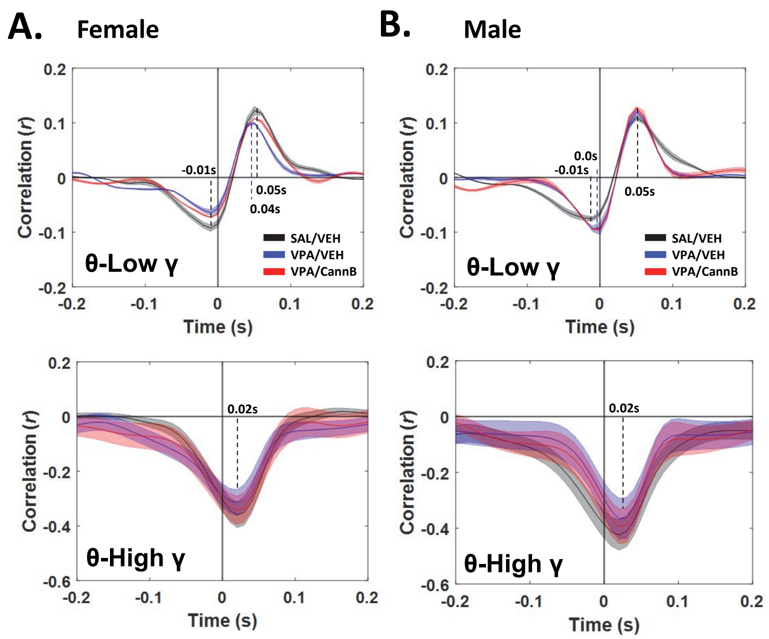


**Supplemental Figure 1. Cross correlograms of the dHIP-Cg theta-gamma coupling. A, B)** Female and male VPA rats displayed a slight 10 ms shift in theta-low gamma coupling that was normalized by cannflavin B in the female, but not male, rats.. N = 10 - 12 animals / group.
